# Comparative metabolomics of wild and cultivated *Taxus cuspidata*: insights into metabolic diversity and environmental adaptability

**DOI:** 10.1101/2024.07.29.605693

**Authors:** Dandan Wang, Yanwen Zhang

## Abstract

*Taxus cuspidata* is a well-known gymnosperm with great ornamental and medicinal value. The study aims to reveal the metabolic differences between wild and cultivated species of *T. cuspidata*, and to analyze the genetic and ecological factors behind these differences. This study conducted a comparative metabolomics analysis of wild and cultivated *T. cuspidata* based on LC-MS/MS technology. The results showed: (1) A total of 7030 metabolites were identified, primarily including flavonoids, organic acids, phenolic acids, amino acids and their derivatives, lipids, and alkaloids, among others; (2) 2381 differential metabolites were confirmed: 949 higher in cultivated species and 1432 higher in wild species, which suggesting wild species have more advantageous metabolites. (3) Through KEGG annotation, 20 significant metabolic pathways were identified, with sugar metabolism, photosynthesis, and sulfur metabolism were significantly higher in wild species. In contrast, pathways related to most amino acids and their derivatives, flavonoids, terpenoids, polyphenols, alkaloids, and plant hormones were significantly higher in cultivated species. In conclusion, the wild species are inferior to cultivated species in terms of drought resistance, growth rate, and medicinal value for cancer treatment. However, the advantage of wild species lies in their significantly higher diversity of upregulated secondary metabolites due to their rich genetic resources, which exhibit greater metabolic diversity and environmental adaptability, which enhances their survival capability. Furthermore, the wild species exhibit significantly higher efficiency in photosynthesis and sugar metabolism compared to cultivated species, enabling them to better withstand low-temperature stress. This study has provided the direction for research into the mechanisms of stress resistance in *T. cuspidata* and has also established a theoretical basis for formulating scientifically and reasonable conservation strategies for wild species.

## 1. Introduction

Plants can be categorized into wild and cultivated population based on significant differences in their growth patterns. Generally, plant populations that grow freely in the wild without direct human intervention and possess the characteristic features of the species model are referred to as wild populations. In contrast, plant populations that have undergone artificial selection, breeding, and improvement, exhibiting significant trait differentiation, are termed cultivated populations (Wang et al. 2017). Wild and cultivated populations exhibit significant differences in morphological characteristics, physiological and biochemical properties, secondary metabolite synthesis, genetic diversity, and other aspects (Roucou et al. 2018). Taking the *T. cuspidata* in Northeast China as an example, under drought stress, the wild species show higher expression levels of key enzymes in sugar metabolism and higher photosynthetic parameters compared to cultivated species. However, cultivated species contain higher content of antioxidant secondary metabolites (Wang et al. 2022). Further analysis reveals that cultivated populations exhibited richer genetic diversity compared to wild populations (Wang et al. 2024). In the study of *T. chinensis*, cultivated species have significantly higher content of most amino acids and flavonoid metabolites compared to wild species, while the cultivated species have lower contents of differential metabolites in sugar metabolism, lipid synthesis, and the TCA cycle pathways compared to wild species. (Chen et al. 2019). Furthermore, similar phenomena are widely observed in other species. For instance, the cultivated species of *Glehnia littoralis*, which exhibits stronger drought tolerance compared to wild varieties (Liu et al. 2023); the cultivated species of *Ganoderma lucidum* exhibits superior antioxidant capacity compared to wild species (Milovanovic et al. 2023); cultivated tomato species emit a more diverse composition of HIPVs than wild species when attacked by psyllids and demonstrating significantly better pest resistance (Bautista-Lozada A et al. 2023). Between wild and cultivated ginseng species, there are a total of 72 significantly different secondary metabolites and 102 differentially expressed proteins. Among these, wild ginseng shows higher biosynthesis and diversity of ginsenosides and phytosterols compared to cultivated species (Ma et al. 2023); The wild banana species have significantly larger average fruit weight, fruit length, fruit circumference, and number of seeds per fruit compared to cultivated populations (Haile et al. 2023); the cultivated species of *Ipomoea batatas* shows lower intraspecific genetic variation due to long-term breeding and domestication, resulting in significant genetic differentiation from wild species (Amritha et al. 2024). In summary, the domestication process from wild to cultivated species is a double-edged sword. On the one hand, it leads to the acquisition or enhancement of certain desirable traits; on the other hand, it inevitably results in the loss or weakening of some potential functional genes. These complex genetic and metabolic variation commonly shapes the significant differences in morphology, physiology, metabolism, and genetics between wild and cultivated populations (Liu et al. 2020).

*T. cuspidata* belongs to the genus *Taxus* in the family Taxaceae, which is a Tertiary relict species. It is suitable for growth in loose, moist, well-aerated, and well-drained soil, and its optimal growth temperature ranges from 20 to 30°C. It is shade-tolerant and cold-resistant, capable of surviving in extreme low temperatures down to -20°C. *T. cuspidata* naturally distributed in northeastern China, Hokkaido and Kyushu in Japan, Sinuiju in North Korea, and islands in the Russian Far East. due to the fact that *T. cuspidata* is the only extremely small population in the northeastern China, with fragmented habitats and discontinuous distribution, it has been listed as a Class I rare and protected plant by the Chinese government (Cheng et al. 2015). It is a valuable tree species with economic, industrial, and medicinal value, which holds a crucial position in the development of anticancer drugs due to its unique bioactive compound, paclitaxel (Zhang et al. 2021). With the deepening of research, scientists have extracted various secondary metabolites from yews, including terpenoids, lignins, flavonoids, alkaloids, and volatile oils, and these discoveries further demonstrate its broad potential for medical applications. However, the current conservation status of wild *T. cuspidata* resources is worrying. On the one hand, the high mortality rate of wild seedlings severely limits the natural regeneration of the population; on the other hand, the surge in market demand for medicinal components has intensified the over-exploitation of wild resources. Facing this dilemma, artificial cultivation of yew has become an important approach to alleviate resource pressure and achieve sustainable utilization. Nonetheless, there are also significant differences between wild and cultivated species in various aspects (Wang et al. 2023). Regrettably, in-depth and systematic comparative studies between wild and cultivated species are relatively scarce at present, which limits our comprehensive understanding of its ecological characteristics, genetic diversity, and medicinal value, while also hinders the formulation of sustainable utilization and conservation strategies.

The liquid chromatography-tandem mass spectrometry (LC-MS/MS) technology, due to its characteristics of high throughput, high resolution, and high sensitivity, is particularly suitable for identifying metabolites that are non-volatile or thermally unstable, which compensates for the limitations of gas chromatography-mass spectrometry (GC-MS) technology in this aspect. When combined with advanced data analysis software such as Progenesis QI v3.0, it can detect and identify hundreds to thousands of metabolites in a single analysis, greatly improving the efficiency and comprehensiveness of metabolomics research (Shi et al. 2024). Metabolomics has demonstrated its remarkable potential in the field of yew research. For example, Zhang et al. (2013) employed GC-MS to analyze the inhibitory substances in the seed coat of *T. mairei*, discovering that the major components include hexadecanoic acid, octadecanoic acid, 9-octadecenoic acid, and 9,12-octadecadienoic acid. Subsequently, Chen et al. (2019) utilized LC-MS technology to reveal the dynamic changes in metabolites such as sugars and flavonoids between wild and cultivated species of *T. wallichiana* var. *mairei*. In the research of taxane compounds, Yu et al. (2020) further confirmed the specific accumulation pattern of taxanes in the phloem of yew stems using metabolomics techniques and elucidated their biosynthetic mechanisms. Gai et al. (2020) employed LC-MS/MS technology to simultaneously detect differences in the content of taxanes and flavonoid compounds in the branches and leaves of three yew species. Recently, Shao et al. (2024) used LC-EI-MS technology to reveal significant differences in secondary metabolites between the heartwood and sapwood of *T. chinensis*, which found that the heartwood is rich in secondary metabolites such as paclitaxel, while the sapwood lacks these key components. Furthermore, Yan et al. (2024) employed metabolomics techniques to analyze the metabolic differences in red, yellow, and purple fruits of *T. mairei* at various developmental stages. However, despite the remarkable progress made in using *T. cuspidata* as a raw material for anticancer drugs, research on the metabolic differences between wild and cultivated species remains relatively scarce. Considering this, the present study aims to fill this gap by focusing on wild and cultivated species of *T. cuspidata* in Northeast China based on LC-MS/MS technology, the study endeavors to:(1) comprehensively detect and compare the metabolite composition of wild and cultivated varieties to clarify the differences in metabolite types and contents between them; (2) Through differential metabolite analysis, explore the differences between wild and cultivated species in terms of stress resistance, adaptability, genetic diversity, growth rate, and medicinal value. Additionally, analyze the significantly different metabolites and their related metabolic pathways; (3) Investigate their metabolic response mechanisms under environmental stresses (such as low temperature, drought, and pest infestations) to provide the theoretical support for the improvement of cultivated species and the conservation of wild species. This study aims to establish a differential metabolite inventory between cultivated and wild species and clarify the advantageous mechanisms of cultivated varieties compared to wild varieties in terms of growth rate, drought resistance, and medicinal value. These findings will not only provide theoretical support for the improvement of cultivated species and the conservation of wild populations, but also offer scientific basis for developing strategies for sustainable utilization and protection of yew resources. Furthermore, this research will promote the application of yew species in the pharmaceutical field.

## 2. Materials and Methods

### 2.1 Experimental Materials and Test Platform

Between 2023 and 2024, the research team selected wild materials (marked as W) from multiple natural distribution areas of *T. cuspidata* in Northeast China,including Liaoning Province (Dandong, Kuandian, Fengcheng, and Tongxing), Jilin Province (Tonghua, Baishan, Fusong, Wangqing, Dunhua, Panshi, Jingyu, Baihe, and Helong), as well as Heilongjiang Province (Muling), a total of 18 representative wild individual samples were selected (Fig. 1, Table 1); For comparative study purposes, 18 cultivated individual samples (marked as C) were collected from multiple large-scale yew plantations, which encompassed a rich diversity of genetic variants, including Japanese yew (*T. cuspidata* var. *nana*), hybrids between Northeast yew (*T. cuspidata*) and Japanese yew, golden yew, and other variant varieties of *T. cuspidata* with unclear origins. For both wild and cultivated types, female plants with similar diameter classes and growth potential were selected. All specimens were identified as *T. cuspidata* species by Professor Yanwen Zhang from Liaodong University. To ensure genetic heterogeneity, the distance between yews was generally greater than 100 meters. The freshly collected leaves were immediately placed in liquid nitrogen for rapid freezing, then transported to the laboratory and stored in a -80°C ultra-low temperature freezer to maintain the physicochemical properties of the samples unchanged, ensuring the accuracy and reliability of subsequent metabolomics analyses. Six biological replicates were set up for each sample to enhance the statistical significance of the data.

**Table 1.** The information on collected wild and cultivated samples of *Taxus cuspidata* from Northeast China.

| Identifier | Admin. region | Altitude<br>(m.a.s.l.) | Tree height (m) | DBH (cm) | location |  |
| --- | --- | --- | --- | --- | --- | --- |
|  |  |  |  |  | Latitude (N) | Longitude (E) |
| W1 | Liaoning Dandong | 113 | 4.2 | 10.8 | 39°44'53" | 123°23'22" |
| W2 | Liaoning Dandong | 121 | 6.7 | 19.2 | 39°44'51" | 123°23'23" |
| W3 | Liaoning Dandong | 129 | 5.1 | 16.7 | 39°44'52" | 123°23'21" |
| W4 | Liaoning Dandong | 129 | 4.8 | 15.1 | 40°53'15" | 124°40'29" |
| W5 | Liaoning Dandong | 129 | 4.6 | 15.6 | 41°10'19" | 124°39'57" |
| W6 | Liaoning Dandong | 129 | 4.9 | 16.9 | 40°24'44" | 124°13'46" |
| W7 | Jilin Baishan | 1415 | 4.2 | 10.3 | 41° 36' 43" | 126° 29' 50" |
| W8 | Jilin Wangqing | 879 | 5.7 | 15.4 | 43°47'44" | 129°48'33" |
| W9 | Jilin Baihe | 846 | 3.8 | 13.1 | 42°24'40" | 128°4'36" |
| W10 | Jilin Jingyu | 953 | 4.1 | 14.6 | 42°10'49" | 126°34'58" |
| W11 | Jilin Gunhua | 799 | 5.9 | 18.3 | 42°42'33" | 127°28'57" |
| W12 | Jilin Helong | 568 | 6.5 | 14.6 | 42°01'03" | 128°47'11" |
| W13 | Heilongjiang Muling | 1256 | 5.1 | 17.6 | 43°50'10" | 129°46'09" |
| W14 | Heilongjiang Muling | 1256 | 5.1 | 18.1 | 44°22'32" | 129°49'50" |
| W15 | Heilongjiang Muling | 1256 | 5.2 | 17.9 | 43°49'41" | 129°50'11" |
| W16 | Heilongjiang Muling | 1256 | 5.3 | 17.3 | 43°51'27" | 130°19'26" |
| W17 | Heilongjiang Muling | 1256 | 5.3 | 18.7 | 44°06'34" | 130°20'21" |
| W18 | Heilongjiang Muling | 1256 | 5.7 | 18.4 | 44°01'43" | 130°28'05" |
| C1 | Liaoning Dandong | 169 | shrub | 4.9 | 40°16'26" | 124°24'42" |
| C2 | Liaoning Dandong | 169 | shrub | 4.7 | 40°37'44" | 124°39'09" |
| C3 | Liaoning Dandong | 169 | shrub | 4.4 | 40°36'19" | 124°41'33" |
| C4 | Liaoning Dandong | 74 | 3.3 | 7.4 | 40°11'11" | 124°15'38" |
| C5 | Liaoning Dandong | 74 | 2.9 | 6.3 | 40°45'27" | 124°06'30" |
| C6 | Liaoning Dandong | 74 | shrub | 5.6 | 41°17'06" | 125°51'41" |
| C7 | Liaoning Dandong | 113 | shrub | 12.6 | 40°14'56" | 124°28'35" |
| C8 | Liaoning Dandong | 113 | shrub | 9.3 | 40°14'35" | 124°26'19" |
| C9 | Liaoning Dandong | 372 | shrub | 12.7 | 40°43'39" | 125°34'20" |
| C10 | Liaoning Dandong | 372 | shrub | 11.1 | 40°46'47" | 123°53'17" |
| C11 | Liaoning Dandong | 372 | shrub | 9.6 | 44°49'08" | 123°33'17" |
| C12 | Liaoning Dandong | 372 | shrub | 8.2 | 41°36'11" | 125°26'38" |
| C13 | Liaoning Dandong | 68 | 4.1 | 12.6 | 40°07'22" | 124°23'49" |
| C14 | Liaoning Dandong | 68 | 3.7 | 10.7 | 40°07'21" | 124°25'26" |
| C15 | Liaoning Dandong | 68 | 3.3 | 9.1 | 40°07'20" | 124°25'26" |
| C16 | Jinlin Changchun | 561 | 3.9 | 9.2 | 44°24'32" | 125°29'23" |
| C17 | Jinlin Changchun | 561 | 3.4 | 8.3 | 44°21'27" | 125°31'22" |
| C18 | Jinlin Baishan | 326 | 3.5 | 8.6 | 43°06'36" | 126°30'06" |

**Figure 1.**
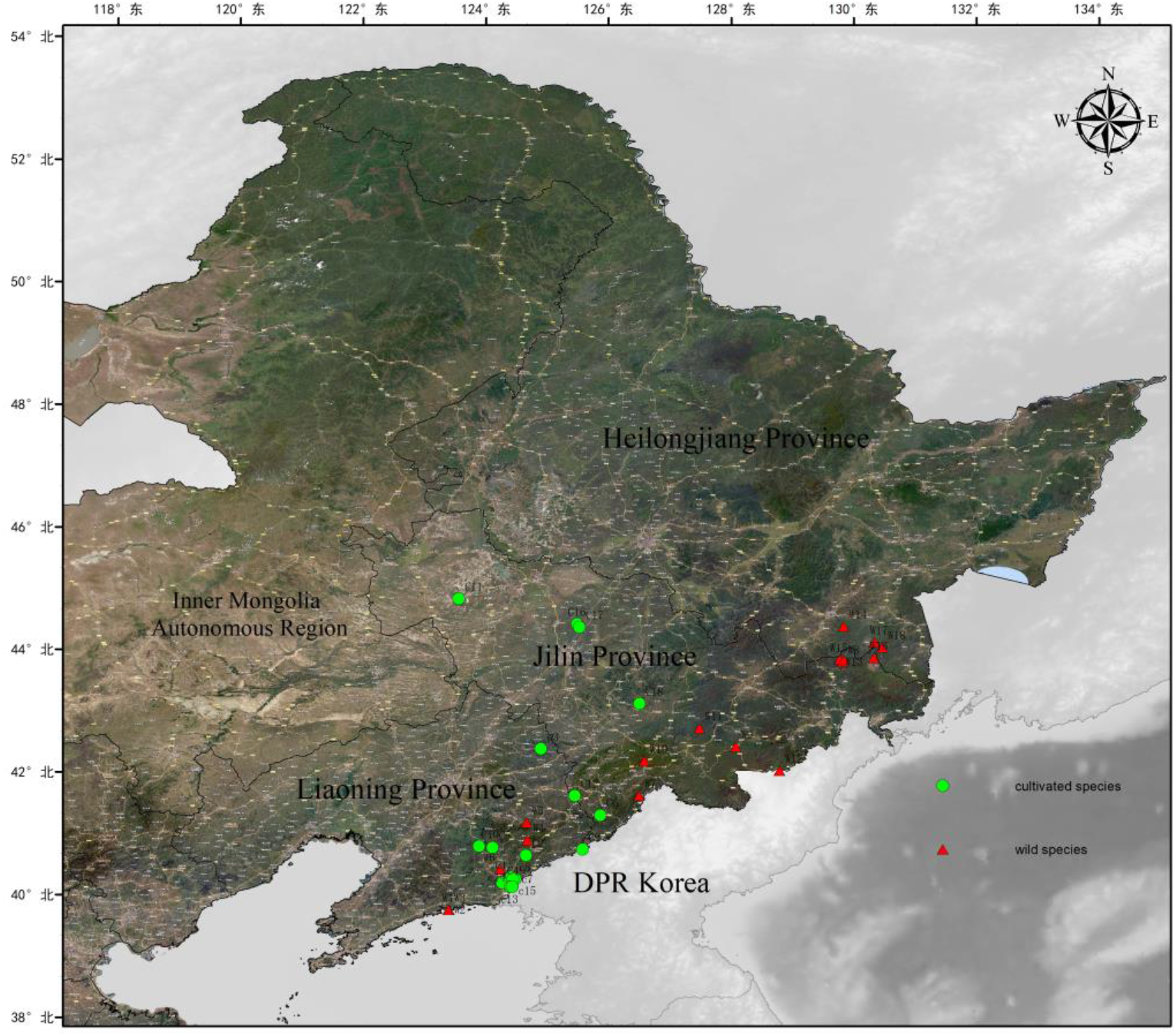
Geographic distribution area of the sampled *T. cuspidata*. The green range represents the sampling locations of the wild species, while the red range represents the sampling locations of the cultivated population.

### 2.2 Extraction of Metabolites from Northeast Yew

Based on LC-MS/MS technology, the research team conducted a comparative metabolomic study between cultivated and wild species of *T. cuspidata* (Dunn et al. 2011; Want et al. 2010). The specific experimental procedures are as follows: Firstly, accurately weigh 60 mg of sample (including both cultivated and wild species) ground in liquid nitrogen and place them into 1.5 mL Eppendorf tubes. Next, add two small steel beads and 600 μL pre-cooled methanol-water mixture (V: V=7:3). After pre-cooling the sample at -40°C for 2 minutes, use a tissue grinder to homogenize the sample (60 Hz, 2 minutes) to further break the cells and release metabolites. Subsequently, the ice-water bath ultrasonic extraction was performed for 30 min to enhance the dissolution of metabolites from the sample matrix. Following this, the sample was kept static at -40°C for 2 hours to promote thorough equilibration and stratification of metabolites with the solvent. Then, centrifugation was conducted for 20 minutes (13,000 rpm, 4°C), and 150 μL of supernatant was aspirated using a syringe and finely filtered through a 0.22 μm organic phase syringe filter before being transferred to LC injection vials. The samples were stored at -80°C until LC-MS/MS analysis. Quality control (QC) samples were prepared by mixing equal volumes of extracts from all experimental samples (Wu et al. 2022). Blank samples (containing only 0.1% formic acid in methanol-water solution) were set up to evaluate experimental background noise and potential chemical contamination. Note: All extraction reagents were pre-cooled at -20°C before use to minimize the impact of temperature changes on metabolite stability, ensuring the accuracy and reproducibility of experimental results.

### 2.3 Analysis using LC-MS/MS Technology

Metabolomics analysis was performed using an advanced integrated liquid chromatography-mass spectrometry (LC-MS/MS) system, combining the Waters ACQUITY UPLC I-Class plus with the Thermo QE ultra-high performance liquid chromatography coupled with high-resolution mass spectrometer. To optimize the chromatographic separation, the ACQUITY UPLC HSS T3 column (100 mm × 2.1 mm, 1.8 μm) was selected, and the column temperature set at 45°C. The analysis was performed in both positive and negative ion modes to comprehensively cover metabolites with different properties. The mobile phase A consisted of 0.1% formic acid in water, and the mobile phase B consisted of 0.35% formic acid in acetonitrile. The flow rate was 0.35 mL/min, and the injection volume for each sample was 3 μL. After every 10 analytical samples, a QC sample was immediately inserted for detection to effectively evaluate the stability of the entire experimental system, ensuring the accuracy and reliability of the obtained metabolomics data.

### 2.4 Multivariate Statistical Analysis of Differential Metabolites

During the data preprocessing stage, ion peaks with a relative standard deviation (RSD) greater than or equal to 0.3 in QC samples were removed to minimize the impact of noise and variation on the analysis results. Data processing and metabolite identification were performed using Progenesis QI v3.0 software (Nonlinear Dynamics, Newcastle, UK), and multiple databases were used for identification analysis, including The Human Metabolome Database (HMDB), Lipidmaps (v2.3), METLIN, and LuMet-Plant3.0, a localized database specifically optimized for plant metabolites. The LuMet-Plant3.0 database covers information on over 10,000 common and specific plant metabolites, broadly encompassing ten major categories of metabolites such as alkaloids, phenolic acids, and flavonoids, and more. During the multivariate statistical analysis stage, an unsupervised PCA method was first used to observe the overall distribution trend and stability between wild and cultivated samples from a macro perspective. Subsequently, to further refine and distinguish the metabolic profile differences between the two groups, supervised statistical methods such as Partial Least Squares Discriminant Analysis (PLS-DA) and Orthogonal Partial Least Squares Discriminant Analysis (OPLS-DA) were introduced.

### 2.5 Data Processing

Based on the comprehensive results of multivariate statistical analysis, key differential metabolites between the two comparative groups were screened. The screening criteria combined the Variable Importance in Projection (VIP) value from the OPLS-DA model, fold change (FC) values, and p-values from Student’s t-test. For this experiment, the screening criteria for significantly differential metabolites were set as VIP > 1, FC ≥ 8.0 or FC ≤ 1/8.0, and p-value < 0.05 (Jiao et al. 2024). To ensure the stability and reproducibility of the experimental results, a 7-fold cross-validation method was used to generate PCA model plots, and a 200-time permutation test was conducted to prevent overfitting of the OPLS-DA model. Comprehensive visualization analysis (volcano plots, box plots, heatmaps, KEGG bubble plots) and correlation analysis of differential metabolites were performed using online tools on the OE Biotech Cloud Platform (https://cloud.oebiotech.com/). Finally, metabolic pathway enrichment analysis was conducted based on the KEGG database (https://www.genome.jp/kegg/).

## 3. Results

### 3.1 Principal Component Analysis

To accurately distinguish the differences in metabolite accumulation between wild and cultivated *T. cuspidata*, SIMCA (V16.0.2) software (V16.0.2) was used for an in-depth analysis of the metabolites in QC samples and the leaves of both types of *T. cuspidata*. An unsupervised PCA model was constructed (Fig. 2). The results showed that all samples fell within the 95% confidence interval, with no abnormal outliers, indicating the good sample quality control and high data reliability. The wild and cultivated samples were clearly clustered and non-overlapping in the PCA plot, clearly indicating significant metabolic differences between the two. Notably, the samples within each group were further subdivided and classified based on their regional distribution differences, for instance, wild samples further subdivided into three clusters based on the three sampling regions (Liaoning, Heilongjiang, and Jilin). it not only reflects the influence of different geographical locations on plant metabolism but also further reinforces the significant metabolic differences between wild and cultivated samples. This multi-level clustering phenomenon not only reveals the fundamental metabolic differences between wild and cultivated yews but also demonstrates the internal consistency of metabolic characteristics within the same ecotype of *T. cuspidata*. It provides important data support for a deeper understanding of the metabolic characteristics and environmental adaptability of *T. cuspidata*.

**Figure 2.**
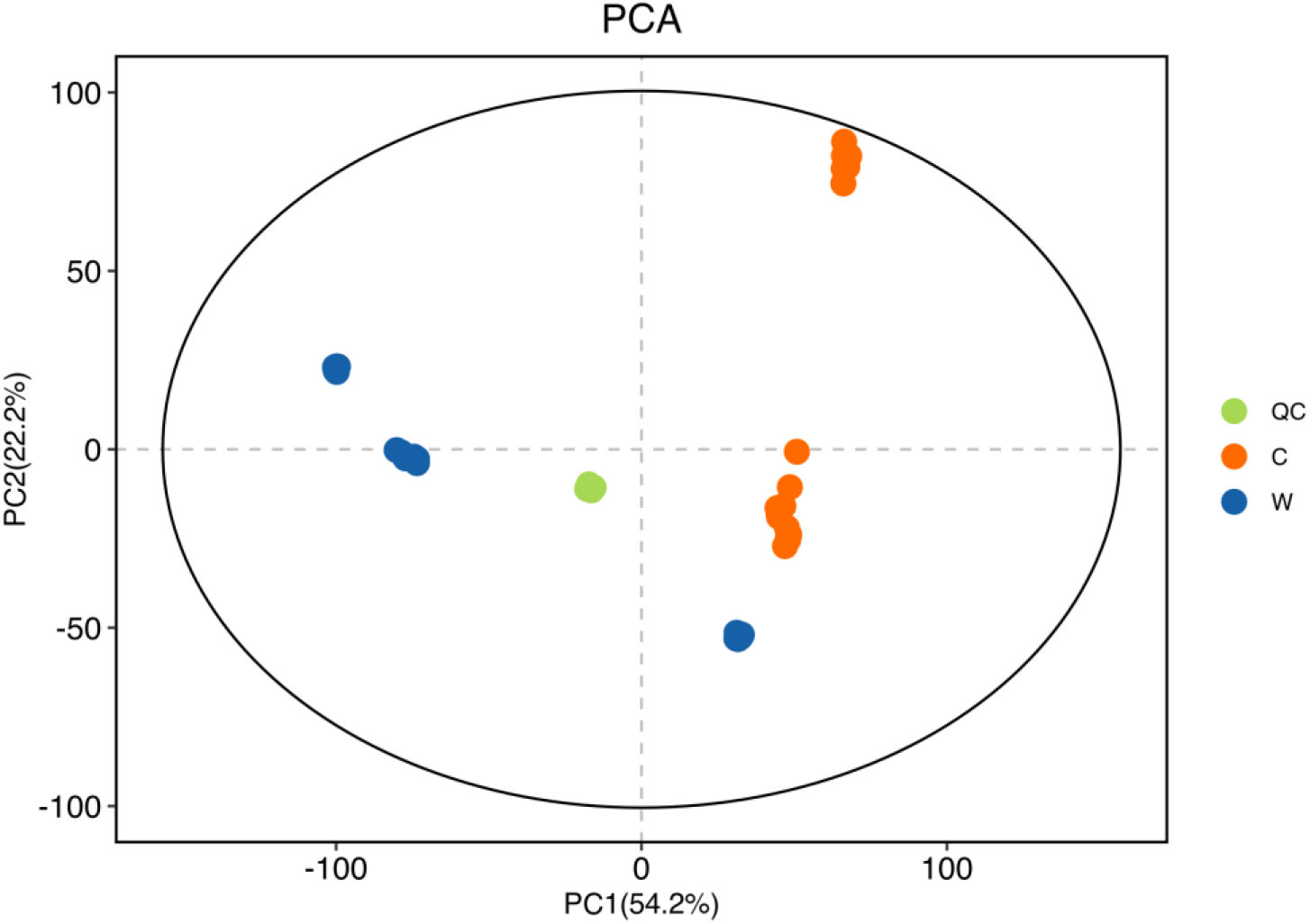
The score plots of PCA for metabolite profiles of wild and cultivated species in *T. cuspidata* analyzed by LC-MS/MS.

### 3.2 Orthogonal projection to latent structures-discriminant analysis

To confirm the metabolic variations between wild and cultivated species, a supervised OPLS-DA model was further employed to optimize group separation, which is an enhancement of PLS-DA, aimed at enhancing intergroup differences. In the 200-response permutation test (Fig. 3B), the key indicator R^2^ had an intercept of 0.139 on the Y-axis, and Q^2^ had an intercept of -0.462 on the Y-axis, both indicators suggest that the OPLS-DA model did not exhibit overfitting and has high reliability. The OPLS-DA results (Fig. 3A) showed significant metabolic differences between the two groups of samples on the first principal component, it not only effectively distinguished between wild and cultivated species but also clearly demonstrated the significant separation characteristics due to different collection sites, which is highly consistent with the PCA analysis. Furthermore, the six biological replicates within each group were highly correlated (correlation coefficients all >0.9), ensuring the repeatability and scientific validity of the data. These results commonly confirm the significant metabolic differences between wild and cultivated species and reflect the influence of different collection sites on plant metabolic characteristics. This provides a reliable data foundation for further in-depth studies on the metabolic characteristics and environmental adaptability of *T. cuspidata*.

**Figure 3.**
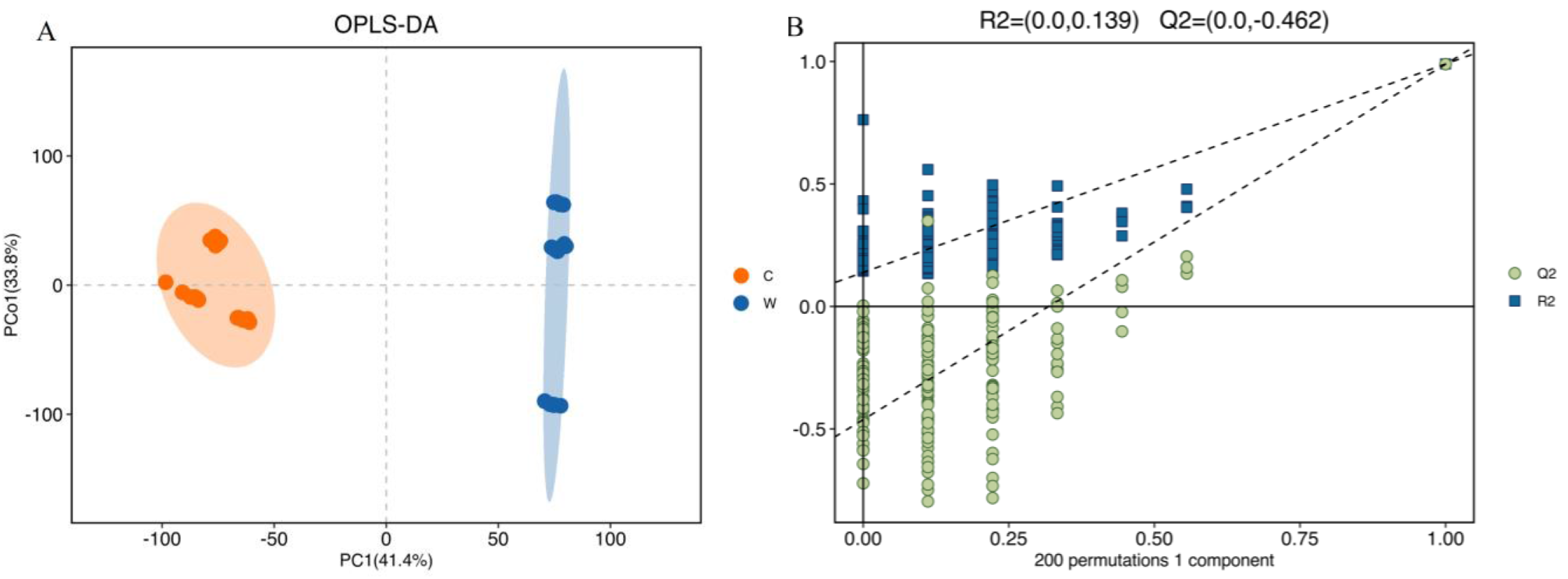
(A) OPLS-DA score plots from metabolite profiles for wild and cultivated samples. (B) A 200 times permutation test of OPLS-DA models for (A).

### 3.2 Analysis of Differential Metabolites between Wild and Cultivated T. cuspidata

In this study, a total of 7030 metabolites were detected between wild and cultivated species. After screening, ultimately confirming 2381 differential metabolites (Fig. 4). These differential metabolites include flavonoids, organic acids, phenolic acids, amino acids and their derivatives, lipids, alkaloids, and others. To control the false discovery rate in multiple comparisons, the Benjamini-Hochberg method was used for multiple testing correction. The screening results were visually presented using a volcano plot (Fig. 5). Compared to the wild species, 949 metabolites were upregulated (marked in red) and 1432 metabolites were downregulated (marked in blue) in the cultivated species.

**Figure 4.**
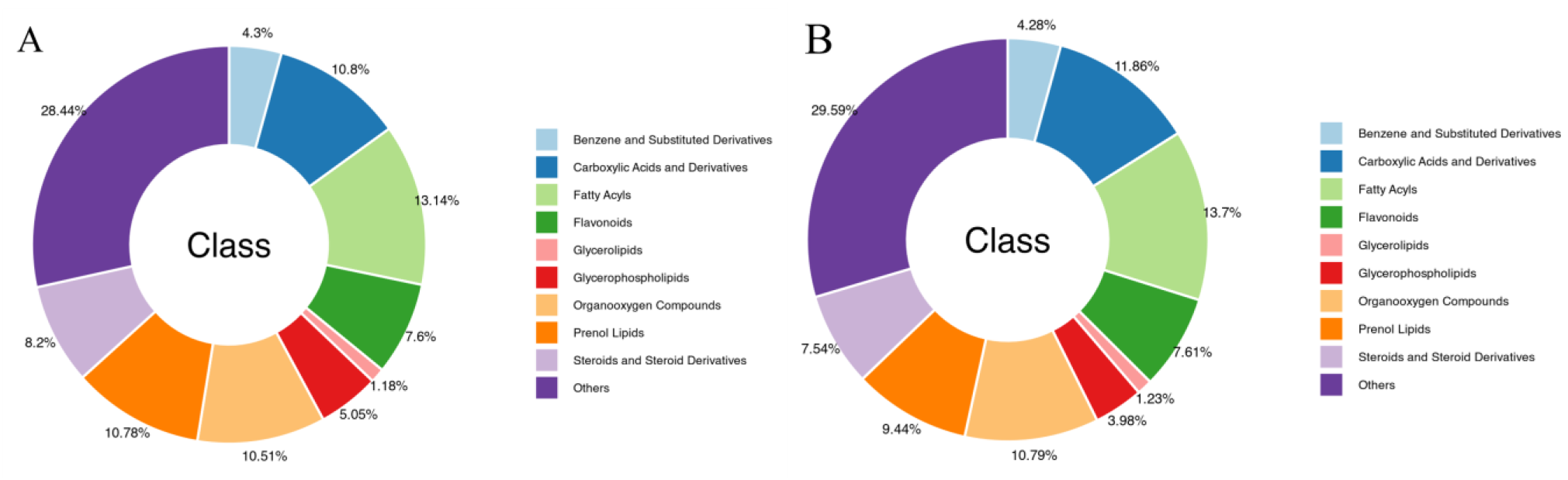
Differential metabolite analysis between wild and cultivated samples of *T. cuspidata*

**Figure 5.**
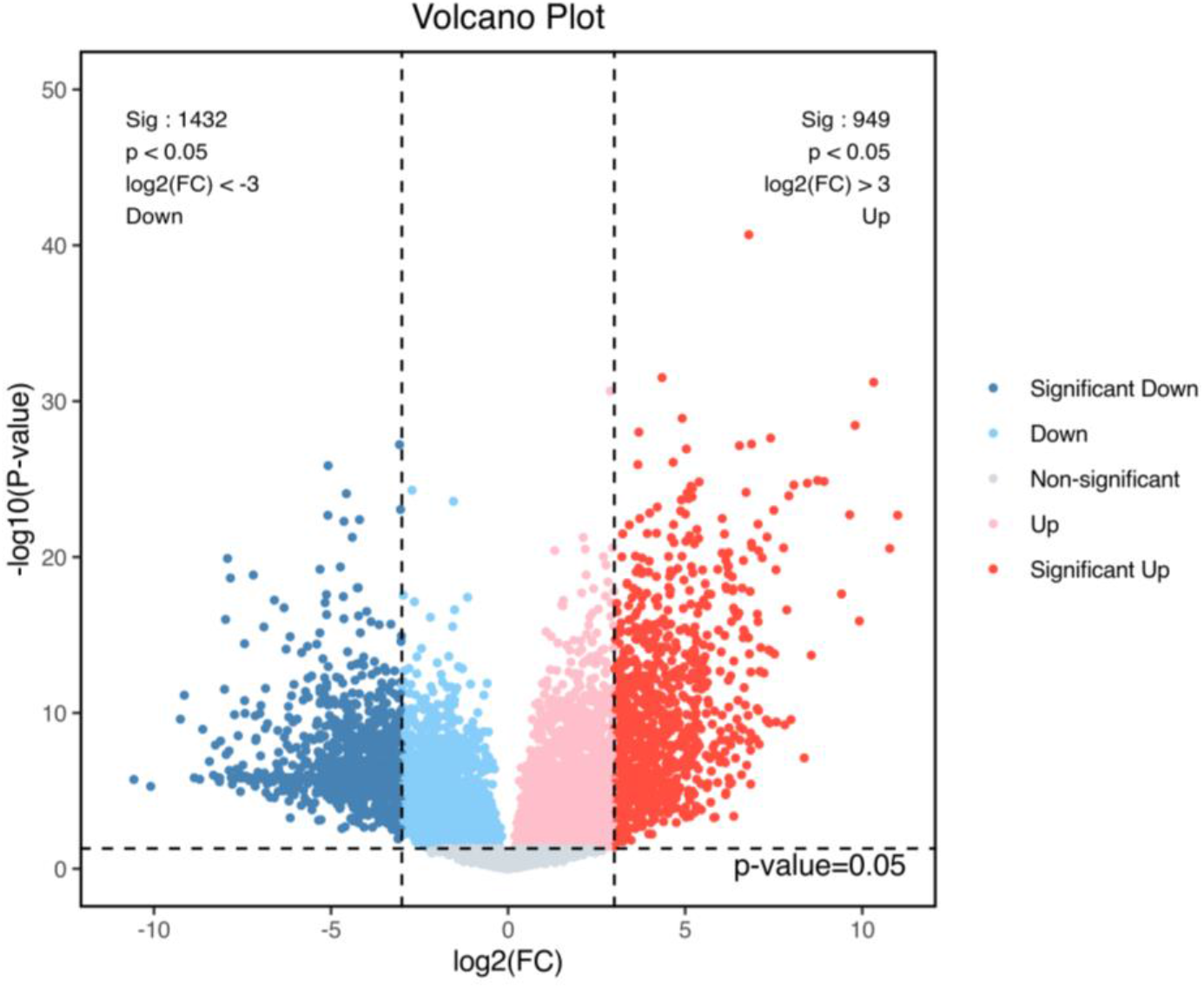
The volcano plot of the differential metabolites between wild and cultivated species of *T. cuspidata*. The red dots represent significantly upregulated differential metabolites, the blue dots represent significantly downregulated differential metabolites, and the gray dots represent non-significant differential metabolites. Each dot in the figure represents a metabolite.

### 3.3 Quantitative Heatmap Analysis of Differential Metabolites

In this study hierarchical clustering analysis was performed on the expression levels of all significantly different metabolites. Based on the minimum p-value, the top 50 significantly different metabolites were selected (Table 2). These metabolites encompass various categories, including fatty acids, phospholipids, alkaloids, aromatic compounds, carbohydrates, vitamins and coenzymes, flavonoids and polyphenols, nucleic acids and nucleotides, plant hormones, amino acids, and peptides. The heatmap (Fig. 6) visually displays the significant differences in metabolite expression abundance between the two, the colors range from blue (low expression) to red (high expression) indicating the relative abundance of metabolites. The clustering results show that the cultivated and wild species formed two distinct clusters, confirming their significant differences in metabolite expression profiles. Compared to the wild species, 31 metabolites were upregulated in the cultivated species, such as PI, DG, 3,4,2’,4’,6’-Pentahydroxychalcone, theaflavin, paclitaxel, tyrosol, glutarylcarnitine, and naringenin; 19 metabolites were downregulated, such as gibberellin A81, tetrahydrobiopterin, spermidine, cyanidin 3-(p-coumaroyl)-glucoside, 3,6-Epoxy-5,5’,6,6’-tetrahydro-β,β-carotene-3’,5,5’,6’-tetrol, cibaric acid, and linusic acid, it indicate that there are significant metabolic differences between the cultivated and wild species, which may impact their biochemical composition physiological characteristics and. In the cultivated species, substances related to sugar metabolism, photosynthesis, and pigmentation are downregulated, while anticancer paclitaxel, most amino acids, fatty acids, phospholipids, phenolic acids, and flavonoids are upregulated, which may be related to artificial selection and optimized cultivation conditions, enhancing the medicinal value, disease resistance, stress tolerance, and growth rate of the cultivated species. Consequently, there are significant differences in the accumulation of primary and secondary metabolites between wild and cultivated species grown under different conditions, demonstrating the key role of metabolites in the growth, development, and stress resistance of *T. cuspidata*. These findings provide important scientific evidence for further optimizing cultivation conditions, enhancing medicinal value, and understanding plant adaptability.

**Table 2.**
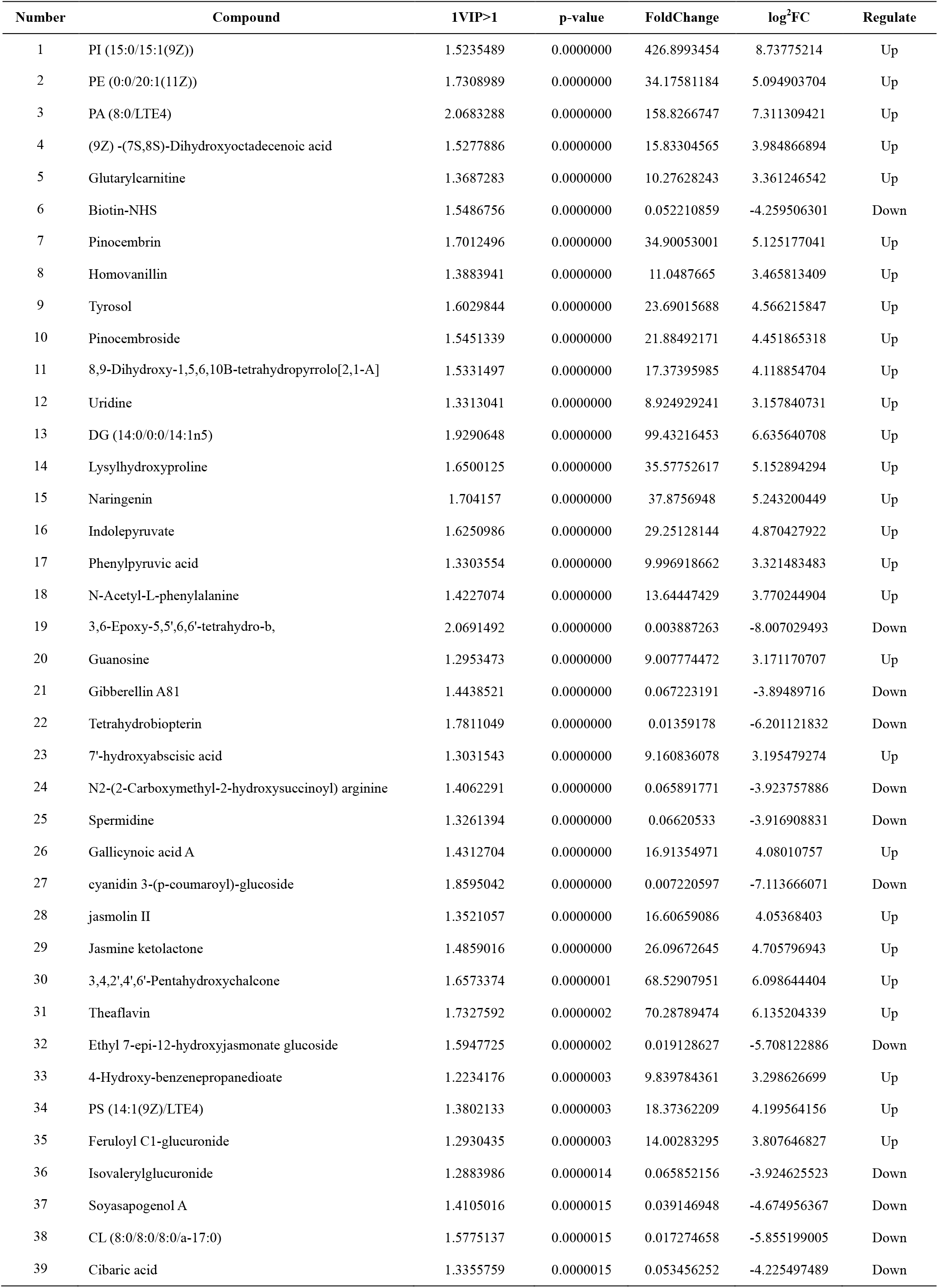
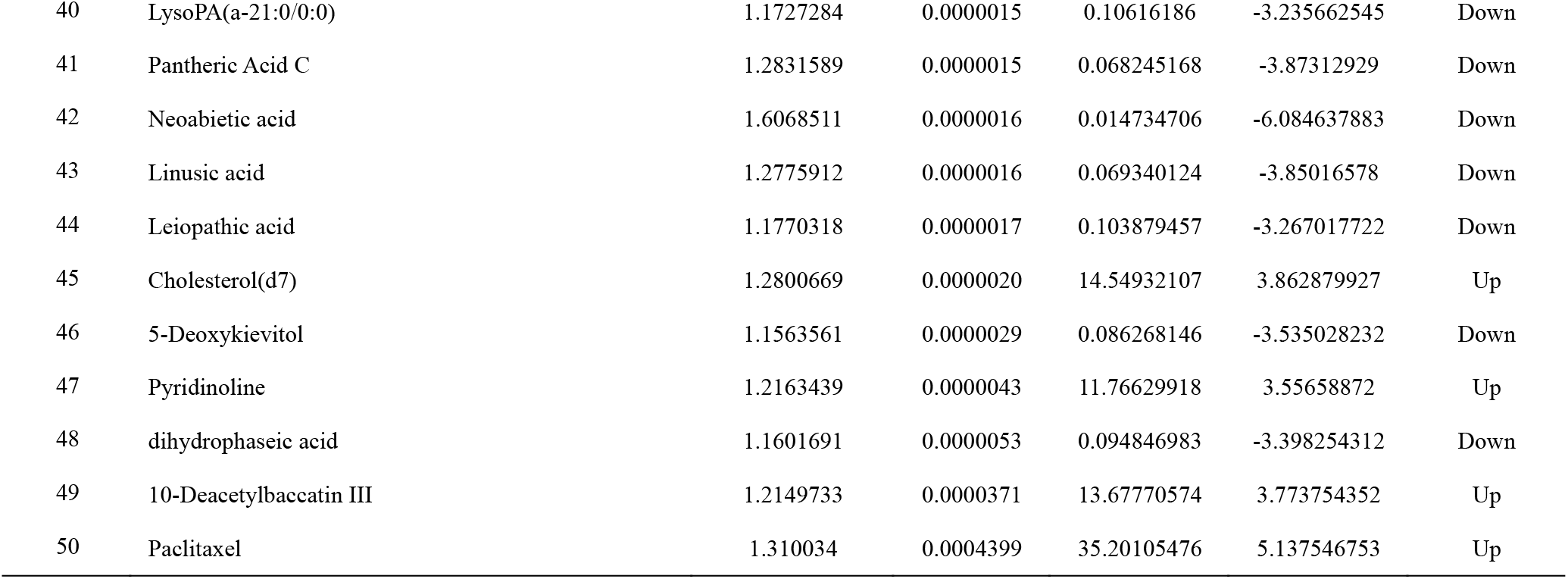
the top 50 significantly different metabolites between wild and cultivated species of *T. cuspidata*.

**Figure 6.**
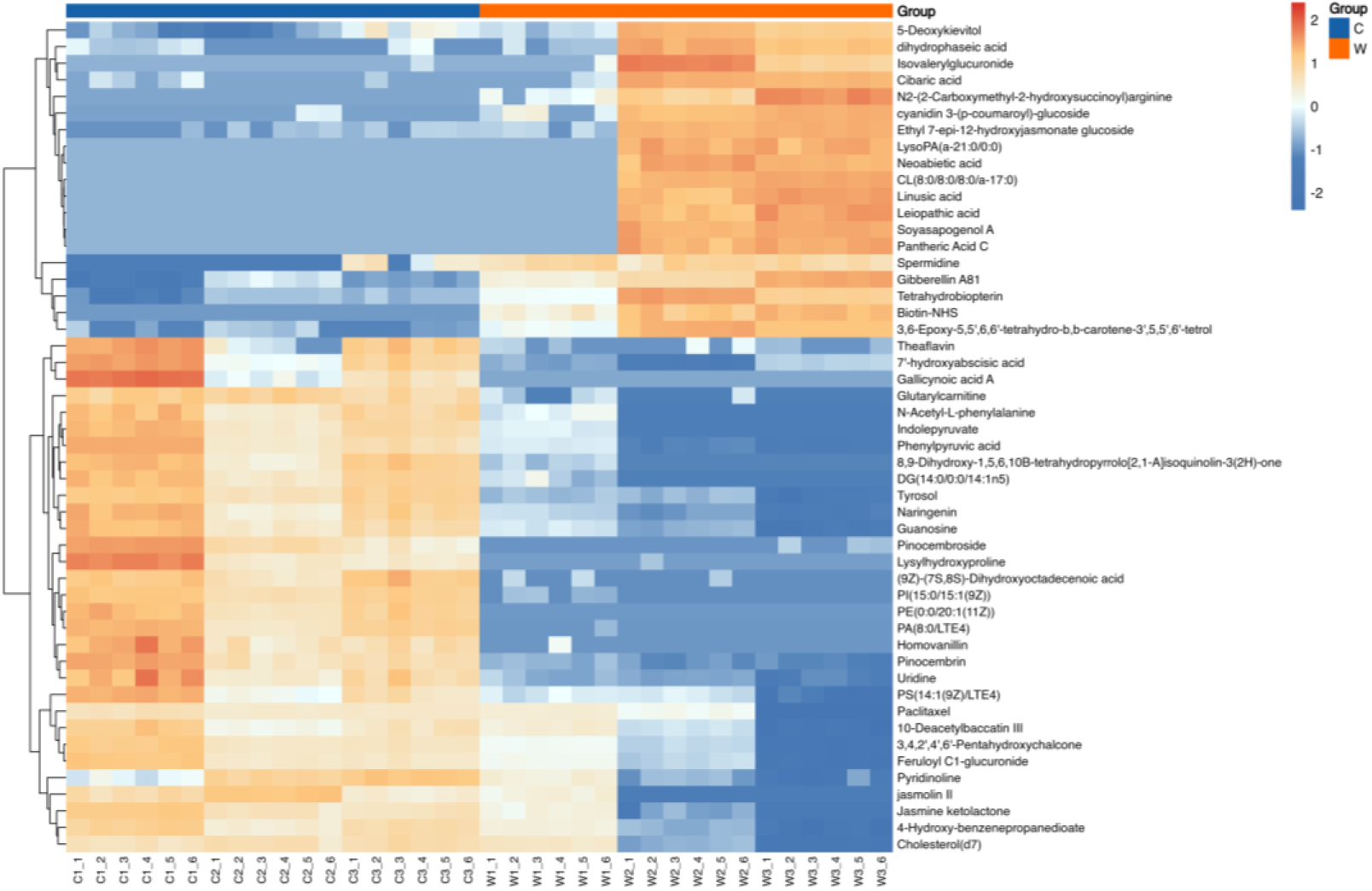
The heatmap analysis of the top 50 differential metabolites between wild and cultivated samples of *T. cuspidata*. The x-axis represents the sample names, and the y-axis represents the differential metabolites. The color gradient from blue to red indicates the expression abundance of the metabolites from low to high, with redder colors indicating higher expression abundance of the differential metabolites.

### 3.4 Metabolic Pathway Analysis of Differential Metabolites

To obtain the pathway information of differential metabolites between wild and cultivated species, metabolic pathway enrichment analysis was conducted using the KEGG database. Based on the minimum p-value, the top 20 statistically significant differential metabolites were selected for further analysis. As shown in Fig. 7A, the significantly upregulated metabolic pathways mainly include cholesterol metabolism, isoflavonoid biosynthesis, pathways in cancer, linoleic acid metabolism, flavonoid biosynthesis, tryptophan metabolism, phenylalanine metabolism, glutathione metabolism, tyrosine metabolism, ascorbate and aldarate metabolism, drug metabolism-cytochrome P450, phenylalanine, tyrosine and tryptophan biosynthesis, lysine biosynthesis, amino sugar and nucleotide sugar metabolism、diterpenoid biosynthesis、glycine, serine and threonine metabolism、steroid hormone biosynthesis、glycerophospholipid metabolism、biosynthesis of various alkaloids、biosynthesis of various plant secondary metabolites, Among these, the metabolic pathways that are significantly upregulated include isoflavonoid biosynthesis、pathways in cancer、flavonoid biosynthesis and tryptophan metabolism. The upregulation of these metabolic pathways in the cultivated species indicates enhancements in antioxidant defense mechanisms, growth and development capabilities, nutrient utilization efficiency, cell signaling, membrane transport functions, anticancer properties, stress resistance, pest and disease resistance, and nutritional value. Consequently, this comprehensive enhancement improves the adaptability and production performance of the cultivated species. As shown in Fig. 7B, the downregulated differential metabolites mainly involve metabolic pathways such as the citrate cycle (TCA cycle), carbon fixation in photosynthetic organisms, beta-Alanine metabolism, lipopolysaccharide biosynthesis, biosynthesis of 12-,14- and 16-membered macrolides, linolenic acid metabolism, pentose and glucuronate interconversions, galactose metabolism, glycolysis/Gluconeogenesis, pyruvate metabolism, amino sugar and nucleotide sugar metabolism, monoterpenoid biosynthesis, cysteine and methionine metabolism, ubiquinone and other terpenoid-quinone biosynthesis, histidine metabolism, biotin metabolism, porphyrin metabolism, sesquiterpenoid and triterpenoid biosynthesis, folate biosynthesis, ABC transporters pathways. Among these, the significantly downregulated pathways include lipopolysaccharide biosynthesis, biosynthesis of 12-, 14- and 16-membered macrolides, and pentose and glucuronate interconversions. Overall, the downregulation of these metabolic pathways in the cultivated species indicates that it is inferior to the wild species in terms of energy metabolism, photosynthesis, and intracellular and extracellular substance transport. This explains the differences between the wild and cultivated species in terms of cold resistance, growth rate, and pigmentation.

**Figure 7.**
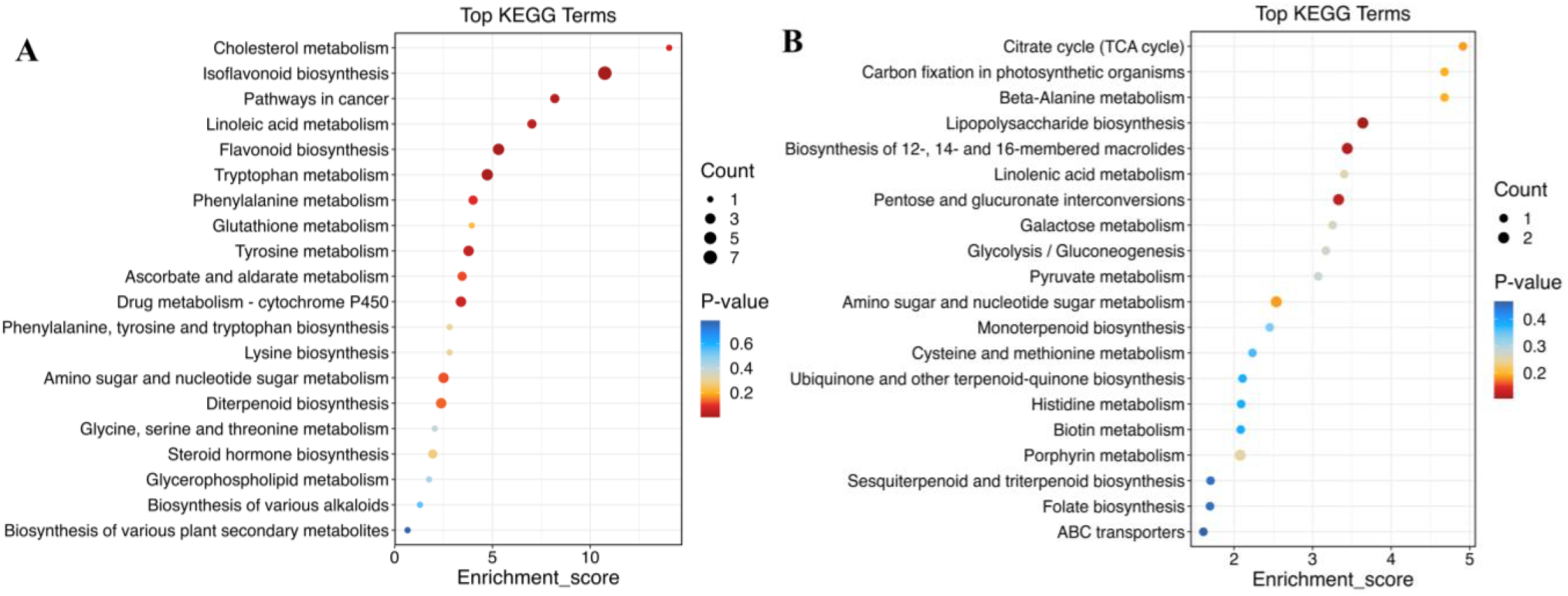
Enrichment analysis of main metabolic pathways of differential metabolites between wild and cultivated samples of *T. cuspidata*. (A) Upregulated pathway of cultivated vs wild species. (B) Downregulated pathway of cultivated vs wild species. The horizontal axis represents the Enrichment Score, and the vertical axis represents the top 20 pathways. (A) (B)The larger the bubble, the more differential metabolites the pathway contains. The bubble color changes from blue to red, indicating that the smaller the enrichment p-value, the greater the significance.

## 4. Discussion

This study, based on metabolomics technology, revealed significant metabolic differences between cultivated and wild species of *T. cuspidata*. These differences may be attributed to the dual effects of their respective living environments and artificial selection practices. Firstly, this study accurately identified the types and quantities of metabolites in both cultivated and wild species. Secondly, it revealed significant differences in stress resistance between the two and elucidated how the wild and cultivated species have evolved unique metabolic strategies to effectively resist environmental pressures in their respective habitats. Thirdly, the differences in metabolite profiles reflect the level of genetic diversity, revealing the mechanisms by which genetic diversity regulates plant phenotypes through metabolic pathways. Fourthly, it analyzed the key metabolites and metabolic pathways that promote the rapid growth and advantageous traits of the cultivated species, providing a scientific basis for breeding high-yield and high-quality cultivated species. Lastly, in terms of medicinal value, it accurately analyzed the changes in the content of medicinal secondary metabolites (such as paclitaxel) in both species, aiming to optimize the cultivated species to enhance their medicinal value, reduce dependence on wild species, and promote sustainable utilization.

### 4.1 Differences in Metabolite Levels Between Wild and Cultivated Species

The wild species significantly surpasses the cultivated species in the following aspects (Figure 6). Firstly, in sugar metabolism, such as the citrate cycle (TCA cycle), pentose and glucuronate interconversions, galactose metabolism, glycolysis and gluconeogenesis, and pyruvate metabolism, which is consistent with the results of Liu’s research (2021). Secondly, in photosynthesis, such as carbon fixation in photosynthetic organisms, folate biosynthesis, porphyrin metabolism, and biotin metabolism, the metabolic levels in the cultivated species are lower than those in wild species. Thirdly, in transmembrane transport proteins, the ABC transporters were more active in wild species. Lastly, in sulfur metabolism, such as cysteine and methionine metabolism. Conversely, the cultivated species demonstrated significantly higher metabolic levels than the wild species in the following aspects. Firstly, in lipid metabolism, such as cholesterol metabolism, linoleic acid metabolism, and glycerophospholipid metabolism. Secondly, in amino acid metabolism, such as tryptophan metabolism, phenylalanine metabolism, tyrosine metabolism, phenylalanine, tyrosine and tryptophan biosynthesis, lysine biosynthesis, and glycine, serine and threonine metabolism. Thirdly, in the biosynthesis of secondary metabolites, such as isoflavonoid biosynthesis, flavonoid biosynthesis, diterpenoid biosynthesis, biosynthesis of various alkaloids, and biosynthesis of various plant secondary metabolites. Fourthly, in drug and toxin metabolism, such as drug metabolism-cytochrome P450. Fifthly, in vitamin and coenzyme metabolism, such as ascorbate and aldarate metabolism. Sixthly, in hormone metabolism, such as steroid hormone biosynthesis. Lastly, in cancer metabolic pathways, such as pathways in cancer. These metabolic differences may be due to long-term selective breeding. During the breeding process, individuals with desirable traits (such as high yield, disease resistance, and superior quality) are often selected for cultivation, leading to the upregulation or downregulation of certain metabolites’ expression, enabling the plants to better adapt to human needs and preferences.

### 4.2 Stress Resistance of Wild and Cultivated Species

The results show that among the top 50 significantly different metabolites identified, most amino acids and organic acids accumulate at higher levels in the cultivated species compared to the wild species. This includes the biosynthesis of unsaturated fatty acids, sulfur metabolism, and the metabolism of arginine, proline, alanine, aspartic acid, and glutamic acid, which help to enhance the plant’s protective mechanisms, energy supply, and cellular repair processes, thereby improving the plant’s ability to withstand various stresses. For example, proline and arginine can effectively maintain cellular osmotic balance and stability under drought stress. Additionally, the accumulation of antioxidants such as flavonoids, polyphenols, alkaloids, and chalcones in the cultivated species far exceeds that in the wild species, and these substances exert strong antioxidant defense capabilities through multiple mechanisms, such as scavenging free radicals, chelating metal ions, and inhibiting oxidase activity, effectively suppressing oxidative stress responses. For instance, the effect of ferulic acid in alleviating drought stress in *Paeonia ostii* also supports this finding (Wang et al. 2020). Furthermore, the content of plant hormones such as abscisic acid (ABA) and jasmonic acid (JA) are significantly upregulated in the cultivated species, which play important defensive roles in stress responses, protection against pathogen and insect attacks. For instance, When ABA (Abscisic Acid) levels rise, it regulates the closure of stomata, reducing water evaporation and enhancing drought resistance. (Zhang et al. 2006). In summary, cultivated species, with their high levels of antioxidant substances and robust stress regulation mechanisms, can respond to and mitigate stress conditions such as drought, pest and disease attacks, and heavy metal pollution more rapidly and effectively. Therefore, it exhibits superior resistance and adaptability to most environmental stresses compared to wild species, and this finding is consistent with research results on *Castanea sativa* (Clark et al. 2023) and *Citrus* spp. (Salonia et al. 2020). From the perspective of photosynthesis and sugar metabolism, the sugar metabolism pathways are crucial regulatory points during low-temperature stress. Low temperatures prompt rapid decomposition of sucrose and starch in wild species, releasing more glucose and fructose for effective energy metabolism to cope with the cold stress. This adaptation may be related to the geographical environment in which the wild species are located, where winter temperatures can reach -20°C to -35°C, or even lower. This finding is consistent with the cold resistance model of blueberries (Rering et al. 2023). Furthermore, the energy (ATP) and organic compounds (such as starch) generated by efficient photosynthesis can regulate the expression of stress-resistant genes, thereby enhancing the plant’s stress resistance (Yang et al. 2022). For example, ATP and NADPH produced during the light reactions provide energy and reducing power for the plant, and NADPH can help eliminate reactive oxygen species (ROS) under stress conditions, reducing cellular damage. Meanwhile, during the Calvin cycle, CO_2_ is efficiently fixed into starch and other organic compounds, which serve as important material bases for stress resistance. This result is consistent with findings in Ugandan coffee (Davis et al. 2023). In summary, wild species rely on efficient sugar metabolism and photosynthetic mechanisms to adapt to extreme low-temperature environments, consistent with the cold resistance models of winter wheat (Fowler et al., 2004) and high-altitude wild tomatoes (Liu et al., 2012).

### 4.3 Genetic Diversity Levels between Wild Species and Cultivated Species

The metabolic levels of *T. cuspidata* in specific ecological environments reflect its genetic diversity and environmental adaptability. Metabolites, as the final products of gene expression and enzyme activity, directly indicate the functional state of genes. Metabolomics analysis can reveal the activation or inhibition state of functional genes in different ecotypes of Northeast yew, reflecting its genetic adaptability characteristics and genetic diversity. For example, under drought stress, cultivated species can accumulate more osmotic regulatory substances (such as proline and soluble sugars) compared to wild species to maintain cellular water balance (Mahmood et al. 2020). Among the 2381 identified differential metabolites, 949 were upregulated and 1432 were downregulated in cultivated species compared to wild species. Although the downregulation content of metabolite was not significant, wild species exhibited stronger environmental adaptability to enhance their survivability, which is consistent with the findings of Zhang et al. (2021). Therefore, by intentionally introducing wild resources from different geographical regions with diverse genetic backgrounds for hybrid breeding, and combining multiple excellent genes, not only has the yield and stress resistance of cultivated species been improved, but their genetic diversity levels have also been significantly increased, as seen in super rice (Zhang et al. 2022). This conclusion is consistent with the findings of Wang et al. (2023). Compared to cultivated species, the numerous upregulated secondary metabolites in wild species indicate they possess rich genetic resources. Although frequent artificial selection typically reduces genetic diversity, scientific breeding strategies and techniques can enhance the desirable traits of cultivated species while also increasing their genetic diversity.

### 4.4 Growth Rates and Morphological Characteristics of Wild and Cultivated species

This study revealed that the growth rate of cultivated species is significantly superior to that of wild species. This advantage is mainly attributed to a series of key metabolites that significantly promote the growth rate of cultivated species on multiple levels, including enhancing metabolic efficiency, strengthening stress resistance, optimizing nutrient absorption and utilization mechanisms, and regulating growth and development processes. Firstly, the upregulation of amino acids associated with growth (like lysine, arginine, glutamic acid, and aspartic acid) in cultivated species has a significant positive effect on their growth rates, leading to larger and thicker leaves. For example, increasing lysine content in maize significantly improves seedling growth rate and seed protein content. Secondly, the increase in phospholipids, such as phosphatidylinositol and phosphatidylethanolamine, promotes cell membrane formation and root growth, leading to enhanced nutrient absorption capacity and more robust plant growth. However, compared to wild species, cultivated species tend to have thinner and smoother bark with softer xylem. This may be because rapidly growing yew tends to allocate more resources to growth and biomass accumulation in a short period, while thick bark requires more energy and resources to form. Third, flavonoids (like naringenin and theaflavin) have strong antioxidant properties, which can reduce oxidative stress, protect cell health, and thereby increase growth rate; fourth, the increase in phosphatidic acid and nucleoside compounds (like uridine and guanosine) not only optimizes the absorption and utilization of nutrients by cultivated species but also promotes the biosynthesis of DNA and RNA, accelerating cell division and further speeding up the growth process, consistent with the views of Testerink et al. (2005). Fifthly, the upregulation of plant hormones and their precursors (such as 7’-hydroxyabscisic acid, jasmonolide, and indole-3-pyruvic acid) regulates the growth and development processes of cultivated species, thereby promoting growth rates at a deeper level. Sixthly, the upregulation of anti-inflammatory and antioxidant substances (such as paclitaxel and dihydroxy-octadecadienoic acid) can enhance the disease resistance of cultivated species, reduce leaf spots, and result in overall stronger plants. Additionally, the downregulation of carotenoids and anthocyanins, along with the upregulation of naringenin and theaflavin, can cause the leaves and fruits of cultivated species to become lighter, yellowish, or overall uneven in color, such as in golden yew. This inference is consistent with reality, whereas the leaves of wild species are usually dark green, which contribute to increased photosynthetic efficiency and adaptation to low-temperature environments. In summary, the synergistic effects of these various metabolites enable cultivated varieties to exhibit significantly faster growth rates and corresponding morphological characteristics compared to wild species.

### 4.5 Medicinal Value of Wild and Cultivated Species

The studies found that the cultivation process significantly impacts the metabolic network of plants, thereby altering their medicinal value. The upregulated metabolites include some active compounds beneficial to human health, such as anticancer drugs, antioxidants, anti-inflammatory substances, and other secondary metabolites. This may be attributed to long-term selective breeding by humans. Examples of these upregulated compounds include anticancer drugs like paclitaxel, antioxidants and anti-inflammatory drugs like pinocembrin, naringenin, and theaflavin, neuroprotective and cardiovascular protective agents like uridine, tyrosol, and feruloyl c1-glucuronide, as well as antibacterial and antiviral drugs like pinocembroside and pasmolin II, and even antidiabetic drugs like phenylpyruvic acid. Conversely, the downregulated metabolites include some that have unique pharmacological effects in wild species but are suppressed or reduced during cultivation. This leads to a weakening of certain medicinal functions in cultivated species. The medicinal metabolites that are significantly higher in wild species compared to cultivated ones include antioxidants, secondary metabolites, sugars and polysaccharides, and some amino acids. These medicinal components, such as tocopheronic acid, spermidine, anthocyanin 3-(p-coumaroyl)-glucoside, neoabietic acid, and dihydrokaempferol, may confer unique medicinal value to wild species, helping them to withstand harsh environmental conditions. In summary, by combining modern agricultural techniques with biotechnology and hybridizing with wild species, the medicinal component content of cultivated species can be enhanced. This not only provides a highly efficient and stable source of pharmaceuticals but also increases their potential applications and economic value in the medical and healthcare fields.

Therefore, the differences in medicinal value between cultivated and wild species not only reflect changes in their metabolites but also that the cultivation method’s potential impact on medicinal efficacy should be considered when selecting and using medicinal materials. The content of important medicinal components in wild species may be lower than that in cultivated varieties, and the possible reasons for this are as follows: Firstly, cultivated varieties are grown in controlled environments where the survival conditions are optimal, favoring the accumulation of medicinal components. In contrast, wild species face variable natural environments with unstable growth conditions, making it difficult to ensure the accumulation of medicinal components. Secondly, cultivated species have undergone artificial selection, leading to higher levels of medicinal components, whereas wild species are subject to natural selection, which prioritizes survival and reproduction, leading to greater variability in medicinal component content. Thirdly, cultivated species grow quickly and accumulate more biomass, which helps in accumulating more medicinal components. On the other hand, wild species often grow in harsh environments, resulting in slower growth rates and less accumulation of medicinal components. Fourthly, the various environmental stresses on cultivated species (like drought, low temperature, pests, and heavy metals, etc) can be artificially controlled, reducing the stress responses, allowing more energy to be used for the accumulation of medicinal components. whereas wild species may allocate more energy towards defense responses against adverse conditions. In summary, due to the advantages of controlled growth conditions, genetic selection, rapid growth, and effective stress management, cultivated species typically accumulate more medicinal components than wild species. This conclusion is consistent with the findings of Yuan et al. (2016).

## 5. Conclusion

This study conducted a comparative metabolomics analysis of wild and cultivated species of *T. cuspidata* from Northeast China using LC-MS/MS technology. A total of 7030 metabolites were identified, primarily including flavonoids, organic acids, phenolic acids, amino acids and their derivatives, lipids, and alkaloids. After screening, 2381 differential metabolites were confirmed, with 949 metabolites having higher content in cultivated species and 1432 metabolites having higher content in wild species. The upregulated metabolites may be related to the enhanced growth, development, and stress resistance of cultivated species, while the downregulated metabolites may be associated with defense mechanisms or secondary metabolic pathways that are no longer needed during cultivation, but they are essential for wild species to adapt to various environmental stresses in natural habitats. For instance, the accumulation ratios of most amino acids and organic acids in cultivated species are higher than those in wild species, and they possess high levels of antioxidants and regulatory substances that alleviate stress, enabling them to quickly mitigate and eliminate damage caused by drought, pests, and heavy metals, which is beneficial for rapid plant growth. The wild species from Northeast China is inferior to the cultivated species in terms of drought resistance, growth rate, and anticancer medicinal value. However, the photosynthesis and sugar metabolism efficiency of wild species are significantly higher than those of cultivated species, allowing them to better withstand low-temperature stress. Furthermore, the advantage of wild species lies in their rich genetic resources and diverse secondary metabolites. Although the content of these metabolites is not significantly higher than that in cultivated species, they demonstrate metabolic diversity and strong environmental adaptability, enabling them to cope with various natural stresses and enhance survival. Additionally, the photosynthesis efficiency of wild species is higher than that of cultivated species, and their expression of carbohydrate substances is higher, indicating a stronger sugar metabolism capability. This allows them to better withstand low-temperature stress, which may be related to the respective living environments of wild and cultivated species from Northeast China.

However, this study still has certain limitations. Although LC-MS/MS technology has high sensitivity and high resolution, its detection range is limited and cannot cover all secondary metabolites, especially some low-abundance or unstable compounds. Secondly, obtaining sufficient and representative samples poses a challenge, particularly for wild species, due to factors such as geographic location, season, and conservation policies. Furthermore, metabolic differences are influenced by both genetic regulation and environmental factors and relying solely on LC-MS/MS technology makes it difficult to fully reveal the effects of gene expression regulation on metabolites. In the future, the sample size will be expanded to increase sample diversity, and multi-omics data, including genomics, transcriptomics, and proteomics, will be integrated for systems biology analysis. This approach will comprehensively elucidate the mechanisms of metabolic differences, providing direction for research on the stress resistance mechanisms of *T. cuspidata* from Northeast China. It will also offer a theoretical foundation for developing scientific and reasonable conservation strategies for wild species, optimizing cultivation techniques, and realizing sustainable resource utilization.

## Supplementary data

The following materials are available in the online version of this article.

**supplementary Figure S1**. Flowchart of metabolome analyses in this study.

**Supplementary Figure S2**. Different levels corresponding to the number of metabolites information in this study.

**Supplementary Figure S3**. Differential metabolite analysis between wild and cultivated samples of *T. cuspidata*.

**Supplementary Figure s4**. Principal component analysis (PCA) plot obtained through 7-fold cross-validation.

**Supplementary Figure s5**. Perform correlation analysis and plotting for QC samples.

**Supplementary Figure S6**. The number of differential substances in each comparison group in the study.

**Supplementary Figure s7**. The volcano plot of the differential metabolites between wild and cultivated species of *T. cuspidata*.

**Supplementary Figure S8**. The heatmap analysis of the top 50 differential metabolites between wild and cultivated species of *T. cuspidata*.

**Supplementary Figure s9**. Enrichment analysis of main metabolic pathways of differential metabolites between wild and cultivated species of *T. cuspidata*.

**Supplementary Table S1**. The information on collected wild and cultivated samples of *T. cuspidata* from Northeast China.

**Supplementary Table S2**. the top 50 significantly different metabolites between wild and cultivated species of *T. cuspidata*.

**Supplementary Data Set 1**. Enrichment analysis of main metabolic pathways of differential metabolites between wild and cultivated samples of *T. cuspidata*. A) Upregulated pathway of cultivated vs wild species.

**Supplementary Data Set 2**. Enrichment analysis of main metabolic pathways of differential metabolites between wild and cultivated samples of *T. cuspidata*. B) Downregulated pathway of cultivated vs wild species.

**Supplementary Data Set 3**. The score plots of PCA for metabolite profiles of wild and cultivated species in *T. cuspidata* analyzed by LC-MS/MS (PCA-score-C-vs-W).

**Supplementary Data Set 4**. The score plots of OPLS-DA for metabolite profiles of wild and cultivated species in *T. cuspidata* analyzed by LC-MS/MS (OPLS-DA-score-C-vs-W).

**Supplementary Data Set 5**. 2381 differential metabolites, 949 metabolites were upregulatedand, 1432 metabolites were downregulated in the cultivated species compared to wild species.

## Acknowledgement

We thank Chunpeng Zhang for drawing the sampling map, and Chang Sui and Xiao Ma for collecting the samples.

## Author Contributions

W-D.D. and Z-Y.W. designed the research. W-D.D. wrote the article and performed the bioinformation analysis. Z-Y.W. revised the manuscript.

## Availability of Data and Materials

All data included in this study are available upon request by contacting W-D.D.

## Ethics Approval

Not applicable.

